# Pial arteries bend with every heartbeat

**DOI:** 10.64898/2026.09.30.755720

**Authors:** Mia Viuf Skøtt, Leif Østergaard, Dmitry D. Postnov

## Abstract

Arterial pulsations are thought to drive perivascular transport by displacing vessel walls radially. In 11 awake mice, all pial arteries also bend with each heartbeat, displacing the wall by three times the pulse-related dilatation per wall, while veins barely move. The bend has a spatial period of about 20 diameters, arrives with the dilatation, at above 1.6 m/s, and dissipates 450 times less energy in the wall.

---

Every heartbeat dilates the cerebral arteries, and this motion is thought to drive fluid transport in the surrounding perivascular spaces [1–4], a route whose failure has been linked to small-vessel disease and dementia [5]. Two related motions have been measured in vivo: the vessel’s dilatation, and the rigid displacement of the surrounding tissue. Models of the perivascular space therefore prescribe an axisymmetric radial motion of the arterial wall [6–10], and the measurement that established perivascular flow in vivo first removes rigid translation of the frame [3]. An artery is a slender tube, and a tube can bend as well as widen. Flexural pulse waves have been observed to travel along human retinal arteries and the carotid [11]. Whether cerebral arteries bend as the pulse wave arrives remains unknown.

We imaged the cortical surface of 11 awake, head-fixed wild-type mice (4 male, 7 female) through a chronic cranial window, using the preparation of our recent blood flow pulsatility study [12]. Wide-field fluorescence of an intravascular dye was recorded at 100 frames per second for 310 s. The heartbeat was detected in each recording’s brightness trace (11.4 Hz [11.2–11.5]; median [interquartile range] across animals unless stated otherwise), and 2656 to 3238 heartbeats per recording were folded onto one averaged cycle of 25 phase bins. In this averaged cycle we separated three motions: the bulk slide of the entire field of view (FOV), fitted on background pixels and removed; each vessel’s change of diameter (dilatation); and the transverse displacement of its centre line (flexural). The two vessel motions were measured along every labelled pial artery (19) and vein (19), from the foot to the peak of the pulse, and compared with scrambled-timing controls of the same beats (Fig. 1a–c; Methods).

**Fig 1.**
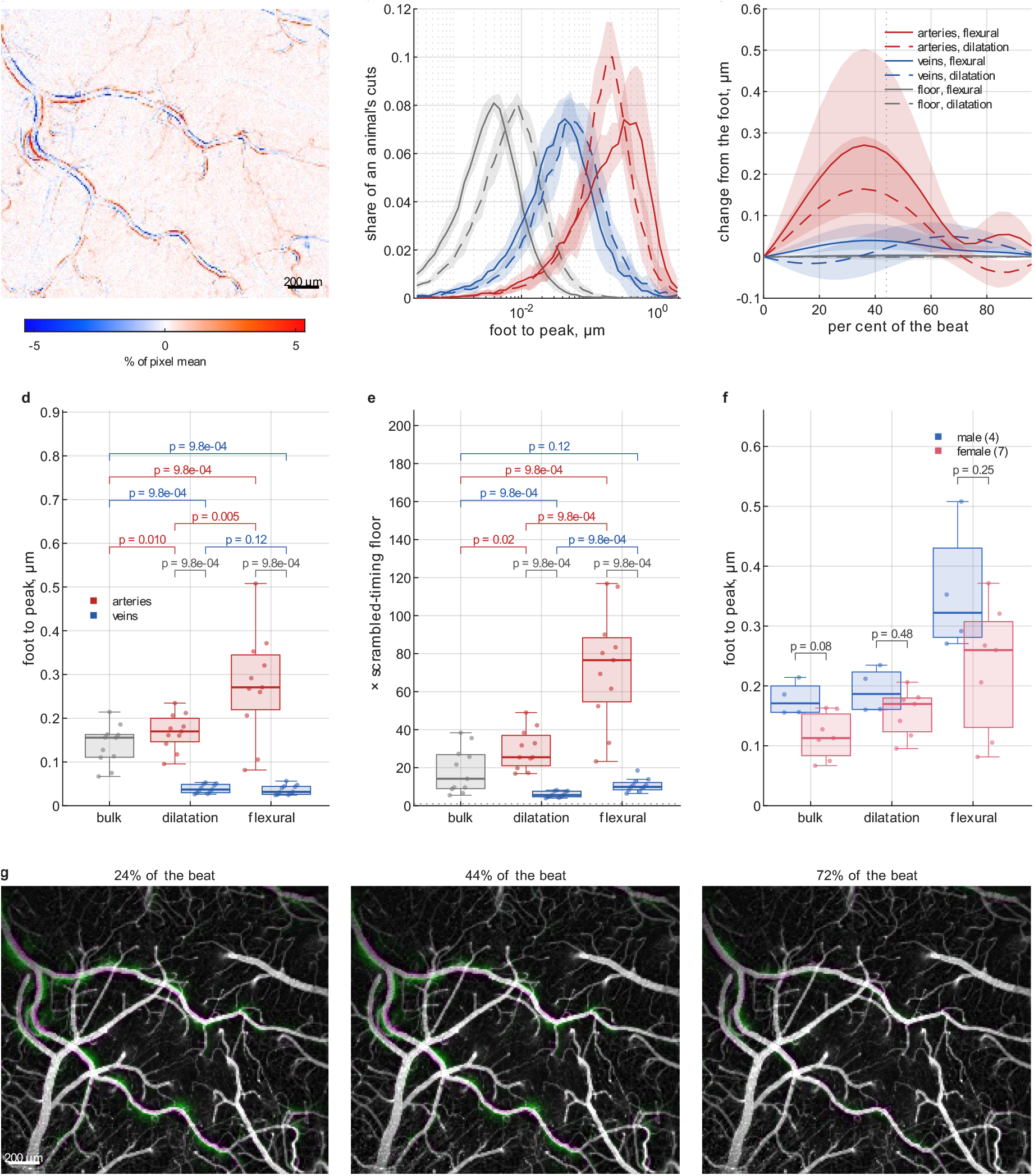
Cardiac-locked motion of cortical vessels. Motion was measured over 39 691 orthogonal cuts placed along automatically segmented arteries (19) and veins (19) in 11 animals. Panels b to f are formed from the cardiac fundamental, and Supplementary Fig. 2 shows them formed from all six fitted harmonics. **a**, Difference in intensity between the peak and the foot of the pulse, for one recording, as a percentage of each pixel’s mean. Red pixels brighten and blue pixels dim (colour limits ±5.35 %). Scale bar, 200 μm. **b**, Histograms of the foot-to-peak change of every cut, for arteries (red), veins (blue) and their scrambled-timing floors (grey). The flexural channel (solid lines) and the dilatation (dashed lines) are drawn as magnitudes on one logarithmic axis. Lines show the mean over the 11 animals and bands ±1 s.d. across animals. **c**, The same cuts over the whole cycle. Lines show the median and bands the quartiles over cuts. The dotted lines mark zero and the peak of the field’s pulse at 44 % of the beat. **d**, Foot-to-peak excursion, with one point per animal (n = 11). Boxes show the median and quartiles, and whiskers reach the furthest point within 1.5 interquartile ranges. Bulk is the slide of the whole field, one number per animal that is not split by vessel type, and dilatation is a change of diameter (not per-wall displacement). Brackets give Wilcoxon signed-rank tests over the 11 animals. **e**, The excursions of d, each divided by the scrambled-timing floor of its channel. The dotted line is at 1. **f**, The excursions of the arteries in male (n = 4, blue) against female (n = 7, red) mice. The first column is the bulk slide of the field. The male medians are larger in all three motions (bulk p = 0.079, dilatation p = 0.48, flexural p = 0.25). **g**, The phase-magnified beat (Supplementary Videos 2 and 3) at 24, 44 and 72 % of the cycle (the peak is at 44 %), for the same recording and crop as in a. Each frame is drawn over the unmagnified foot frame as a false-colour pair, with the foot in magenta and the magnified frame in green. A displaced wall appears as a paired magenta and green fringe. Scale bar, 200 μm.

Arteries bend, and they bend in every animal. The per-animal median of the flexural excursion was 0.271 μm [0.220–0.345]. The pulse-related dilatation was 0.170 μm [0.146–0.200], a change of diameter and so 0.085 μm per wall, and the bulk slide of the FOV was 0.156 μm [0.111–0.163] (Fig. 1d). Per artery, the bend was 3.11 [2.32–4.13] times the dilatation per wall. Across animals it was larger than the full change of diameter as well (p = 0.0049, Wilcoxon signed rank, n = 11). The scrambled-timing floor is what the same analysis returns once the timing between beats is destroyed, and so measures the noise. The bend stood 77 [55–88] times above its floor, the dilatation 25 [21–37] and the bulk slide 14 [9–27] (Fig. 1e). The bend varies widely between arteries, from 0.062 to 0.697 μm (per-artery medians, n = 19). It is often visible to the naked eye in unprocessed frames played ten times slower (Supplementary Video 1). A similar bending can be seen, though it was not reported, in Supplementary Movie 2 of ref. 3, recorded in anaesthetised mice. In veins, measured on the same frames by the same fit, the flexural excursion was 0.032 μm [0.026–0.044] and the dilatation 0.037 μm [0.030–0.049]. Both were five to nine times below the arteries in every animal (p = 9.8 ×10^−4^) and did not differ from each other (p = 0.12). The bend is therefore an arterial motion. The male medians were larger in all three motions, but no difference reached significance (Fig. 1f).

Binned by direction, the flexural excursion is 0.27 μm over most of the circle, with one lobe that the veins do not share (Fig. 2a). To examine whether the motions travel along the artery, we measured the delay from its origin to its far end, with a sign-blind estimator and the camera’s readout delay taken out (Methods). Both motions reach the far end a little after the origin, the flexural wave by 1.94 ms (median across animals, positive in 10 of 11, sign test p = 0.012) and the dilatation by 2.04 ms (10 of 11, p = 0.012). The two lags agree across arteries (r = 0.63, p = 0.0038; Fig. 2b). The lag is below one frame (10 ms), so through-origin fits give only lower bounds on the arrival speed, 1.6 m/s for the flexural wave and 1.1 m/s for the dilatation. To our knowledge no pulse-wave velocity has been previously reported for a mouse pial artery, making our results the first experimental bounds that are also close to the 1 m/s that models of perivascular flow assume for the cardiac wave [10]. Phase magnification makes the bend directly visible as paired magenta and green fringes along the wall (Fig. 1g; Supplementary Video 2; control, Supplementary Video 3). To examine how the bend is laid out along an artery, we mapped the sign of its cardiac fundamental and found that it alternates (Fig. 2c), with nodes that do not move. The spatial period of the alternation, the bending wavelength, was 620 μm [504–803], or 19.7 [14.6–22.9] lumen diameters. It rose with diameter (r = 0.46, p = 0.047), not with the length measured (r = 0.12, p = 0.62).

**Fig 2.**
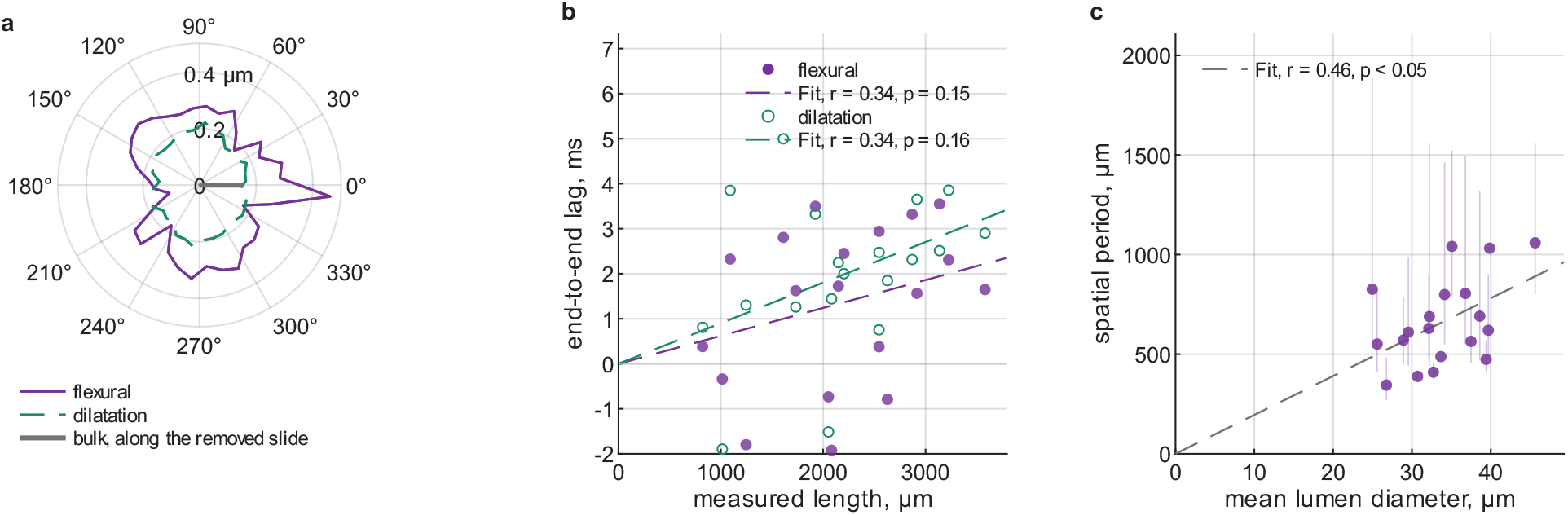
How the motion is laid out along the artery. All panels show the 19 arteries of the 11 animals. Panel a is formed from the cardiac fundamental, and Supplementary Fig. 3 shows its six-harmonic version. Panels b and c are defined on the phase of the fundamental, so they are the same for the six-harmonic version. **a**, Direction of the foot-to-peak motion of the arterial cuts, in the frame of the removed bulk slide. The curves give the mean excursion per direction in 36 bins of 10^*°*^. The flexural motion (solid line) is assigned to the end of the cut’s normal towards which the centre line moves from foot to peak, and the dilatation (dashed line) to both ends. The curve of the dilatation is therefore symmetric under a rotation of 180^*°*^ by construction. The grey line at 0^*°*^ is the removed bulk slide, drawn along its direction at its median length of 0.156 μm. The outermost tick is at 0.4 μm. The flexural curve is largest in the bin next to the direction of the slide (0.46 μm) and smallest opposite it (0.11 μm), against 0.27 μm over most of the circle. The same lobe, however, is not present for veins. We therefore associate it with arterial motion rather than a residue of the removed field motion. **b**, End-to-end arrival lag of the flexural wave (filled circles) and of the dilatation (open circles) of each artery against the length of the measured stretch, with one point per artery (n = 19). A positive lag means a delay while propagating downstream. Its per-artery uncertainty is about 5.3 ms, so the test is of the sign of the lag over animals. The flexural lag is +1.94 ms (median over animals) and is positive in 10 of 11 animals (sign test, p = 0.012). The lag of the dilatation is +2.04 ms and is positive in 10 of 11 (p = 0.012), and the two lags correlate over arteries (r = 0.63, p = 0.0038). The dashed lines are through-origin fits, and their slopes give lower bounds on the arrival speed of 1.6 m/s for the flexural wave and 1.1 m/s for the dilatation. **c**, Spatial period of the signed transverse displacement along each artery, which is the bending wavelength, against the mean lumen diameter of the artery, with one point per artery (n = 19). Whiskers show the confidence interval of the coherence plateau. The dashed line is a through-origin fit at 19.5 diameters. The median period is 620 μm (interquartile range 504–803 μm), or 19.7 [14.6–22.9] diameters. The period rises with the diameter (r = 0.46, p = 0.047) and not with the length measured (r = 0.12, p = 0.62).

We calculated how the energy of both motions dissipates into the wall itself, the perivascular fluid and the brain outside it, using a quasi-steady dissipation model (Methods). The dilatation strains the wall in hoop by 0.256 % [0.236–0.315] and the bend in axial bending by 0.0172 % [0.0108–0.0300], 13.0 [8.0–25.5] times less per artery (Fig. 3a). A viscoelastic loss goes as the square of a strain, with a geometric factor of 8/3. The wall therefore dissipates 448 [172–1761] times more of the dilatation’s energy than of the bend’s, 538 fJ [376–709] per beat and millimetre of artery against 0.70 fJ [0.28–3.22] (Fig. 3b). Outside the wall the answer depends on whether the outer wall of the perivascular space is held or free, which we cannot measure. In the phase-magnified videos the bending arteries appear to drag the surrounding tissue and veins with them, which favours the free limit (Supplementary Video 2). With the outer wall held, the dilatation must vent fluid along the annulus to the nearest opening, while the bend is relieved round the annulus and delivers 29 [17–63] times less to the fluid per artery. With it free, the brain takes the motion and the bend delivers 3.4 [2.2–6.4] times more to it than the dilatation. That one assumption swings the comparison outside the wall by a factor of about a hundred. In the held limit, only the energy the dilatation delivers to the fluid grows with the venting length, and the two are equal at 236 μm [163–310], inside the 50–300 μm spacing of the penetrating arterioles [13, 14] and five times below the 1280 μm bifurcation spacing of these arteries. In the free limit, no loss depends on that length and the bend delivers more at every length (Fig. 3c). If the bend is a helix that turns out of the imaging plane, which a one-plane camera cannot exclude, every flexural energy doubles exactly and that equality moves to 334 μm [230–439].

**Fig 3.**
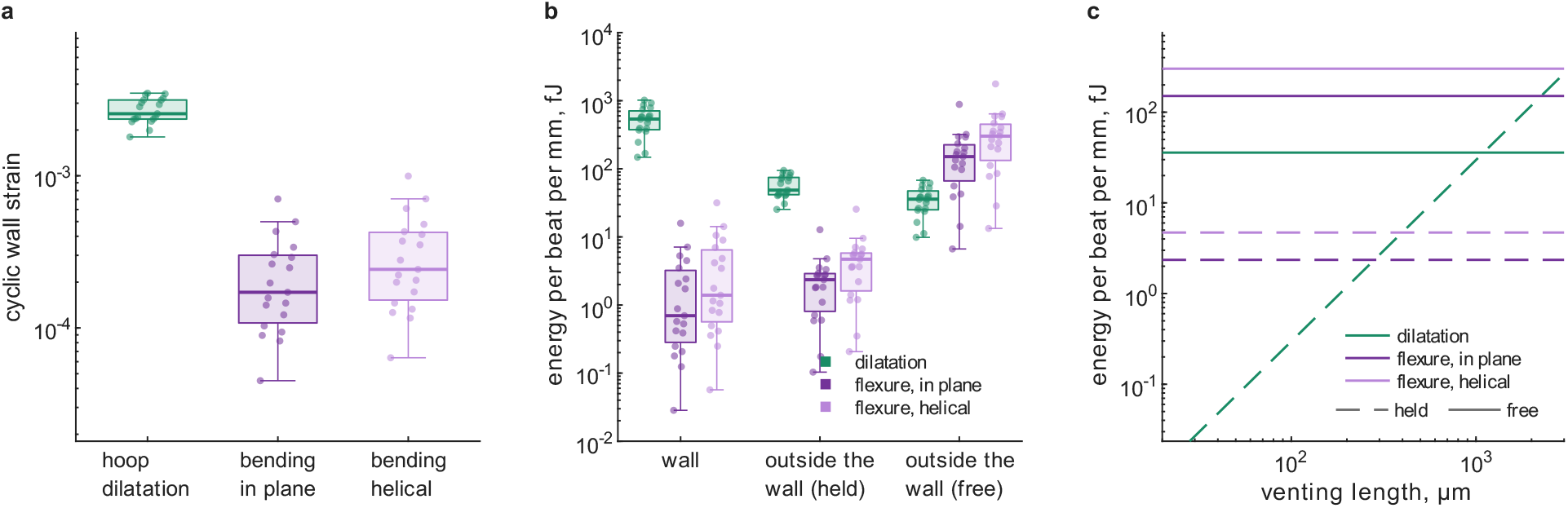
Where the energy of a beat goes, according to a dissipation model. All panels are formed from the cardiac fundamental, and Supplementary Fig. 4 shows the six-harmonic version. Green is the dilatation, purple the bend read in the imaging plane and lighter purple the same bend read as a helix that turns out of the plane. The helical reading doubles every flexural energy exactly, and it is a second reading of one measurement and not a second channel. Each point is one artery, boxes show the median and the quartiles, and the axes are logarithmic. **a**, The cyclic strains in the wall. The dilatation imposes the hoop strain *A/R*_1_ and the flexure imposes the bending strain 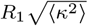, which is shown in the plane and as a helix. The medians are 0.256 %, 0.0172 % and 0.0243 %, and the ratio of hoop to bending strain is 13.0 [8.0–25.5] per artery. **b**, Energy per beat per millimetre of artery, with one box per motion and where it goes. The wall column is the viscoelastic loss of the arterial wall itself. The two outer columns are one channel at its two limits. In the held limit the outer wall of the perivascular space is held, the gap takes the whole excursion and the loss is that of the fluid (assuming a venting length of 1280 μm). In the free limit that wall is free and the loss is that of the brain tissue, which takes the motion where it happens. No measurement chooses between the two limits, so both are drawn, and the comparison between the two motions is likely to fall in between. For the dilatation, the planar bend and the helical bend, the medians in fJ per beat per mm are 538, 0.70 and 1.39 in the wall, 48.6, 2.4 and 4.7 in the held limit, and 35.8, 151 and 302 in the free limit. The flexural wall loss goes as the fourth power of the bending wavenumber, so its box reflects the spread of the bending wavelength (Fig. 2c). **c**, The same channel at both limits against the venting length of the perivascular space, which is the distance between the places where the space opens into another. Every line is a median over the arteries, the spreads are the boxes of panel b, and the two limits are never summed. Dashed lines show the held limit, in which the fluid loss of the dilatation rises as the square of the venting length. That of the bend is flat, because its fluid is relieved locally and never reaches a vent. Solid lines show the free limit, in which no loss grows with the venting length (35.8, 151 and 302 fJ per beat per mm) and the bend delivers more than the dilatation at every length. In the held limit the curve of the dilatation crosses that of each bend where the two fluid dissipations are equal (parity), at 236 μm [163–310] in the plane and 334 μm [230–439] as a helix. Parity is an equality of two dissipations and not a transport threshold, and the tethering cancels out of it because both fluid losses carry *τ* ^2^. The crossings lie inside the 50–300 μm spacing of penetrating arterioles [13, 14] and below the 1280 μm pial bifurcation spacing of this set, at which the held column of panel b is evaluated. Constants and their sources are given in Supplementary Table 1.

The objections to perivascular pumping concern phase and cannot be answered here. A transverse shift of a cylinder inside a larger one preserves the area between them exactly, so if the perivascular space keeps its cross-section, the bend alone would drive no net axial flow at leading order, as the one simulation to translate an artery rigidly also found [15]. It would instead drive an oscillatory flow alternating in azimuth along the vessel, with the mixing and dispersion of solute that a non-axisymmetric geometry is already predicted to produce [8, 16]. But the bend does not act alone. It rides on the same beat as the dilatation, arrives with it and moves the wall three times further per wall, so the arterial wall works on the perivascular space in two ways at once. It changes the boundary condition every model has used. The wall moves in a way that is not axisymmetric, by more than it dilates per wall, and with a layout along the vessel that no model we know of represents. Models of perivascular pumping have consistently predicted a much stronger oscillatory component than is observed [16], a discrepancy that realistic end conditions largely remove [17]. Whether the motion prescribed also matters, and what the bend does for glymphatic transport, is for models that carry it to decide.

Three limits set what this measurement can decide. The arteries measured lie on the cortical surface, in fluid-filled spaces bounded by the pia and the arachnoid [18], and how these tether them was not measured. Whether the bend is planar or helical is undecided. A helical bend leaves the focal plane and should modulate the vessel’s width and brightness where its in-plane displacement is smallest. Finally, whether the bend continues down the penetrating arterioles needs depth-resolved optical imaging along the vessel at sufficient temporal resolution.

## Methods

### Animals and preparation

All procedures were approved by the Danish National Animal Experiments Inspectorate (permit no. 2023-15-0201-01399). Eleven wild-type C57BL/6JRj mice (Janvier Labs; 4 male, 7 female), 12 months old at recording, received a head plate and a chronic cranial window over the left barrel cortex (craniectomy 1.5–2 mm posterior and 3 mm lateral to bregma, a 4 mm round coverslip on Kwik-Sil, sealed with dental cement). Surgery was followed by at least ten days of recovery and ten days of habituation to head-fixed restraint, as described for our pulsatility study [12]. On the day of recording the mouse was lightly anaesthetised for 30–40 min in all, with about 1.5 % isoflurane for the insertion of a tail-vein catheter and about 1 % for cleaning the window. It recovered for at least 20 min before a fluorescein bolus and angiogram, and for a further 10 min before the recording used here. That recording therefore began at least 30 min after the anaesthesia ended, with the mouse awake and head-fixed. Afterwards the catheter was removed under less than 2 min of 3 % isoflurane, and the mouse moved within 30–60 s of returning to its cage. Fourteen recordings were made, and three were excluded before analysis for low cranial window quality.

### Imaging

The dye was an intravenous bolus of 75 μL of 2 % (w/v) FITC-dextran (70 kDa) in saline, given through the tail-vein catheter. Wide-field one-photon fluorescence of the cortical surface was recorded on a Hamamatsu ORCA-Flash4.0 V3 sCMOS camera (C13440-20C) using a 2.5 ×/0.07 objective (Leica N Plan) and a 200 mm tube lens, with LED excitation through a FITC filter set. The camera recorded a sub-array of 1300 lines by 800 columns of the 2048 ×2048 sensor (binning 1, 16 bit, exposure 10 ms). The sub-array was placed on the sensor per animal (column offset 580–780, line offset 348–500). Each recording is approximately 31 000 frames over 310 s at about 100 frames per second. The image scale is 2.65 μm per pixel, measured directly under the objective, so the frame covers 3.4 ×2.1 mm. The field of view (FOV) on which all motions are measured is a cropped rectangle of 1034 lines by 800 columns, placed per recording to hold every labelled vessel. The camera’s standard readout runs from the sensor’s centre line outward to both edges at 9.744 μs per line (manufacturer’s manual A3410301-09, section 11-1). Its consequences for the phase along the vessel are treated below.

### The averaged cardiac cycle

The heartbeat is found in the recording itself. The frame is grouped into 8-pixel blocks, and the brightness trace of the bright (vascular) 15 % of the mean image is detrended. The rhythm at which the two halves of the field agree is then sought between 8 and 15 Hz, in a tournament that separates the heart from breathing and slower waves.

Each beat in that band is accepted or rejected by rules on its duration, prominence, level and shape (2656 to 3238 accepted per recording) and is placed on one cycle by its phase. The cycle is drawn on 25 phase bins, and one harmonic series of up to six harmonics is fitted per pixel. All six were kept in every recording. The main figures formed from all six harmonics are Supplementary Figs. 2 to 4. The five extra harmonics carry little of the vessel motion and much of the noise of the control, so those figures lie closer to their floors. They serve as a check on the fundamental, and nothing is claimed from them that the fundamental does not also support.

Each averaged cycle is accompanied by three scrambled-timing controls. A control is built from the same frames, beats and fit, but each beat is given one random shift around the cycle before averaging. The shift destroys the timing between beats and keeps the tissue, the noise, the exposure and the number of beats. Every quantity below is compared with the median of its three controls, and a control is always fitted, filtered and warped by its own estimate. The averaged cycles of the recordings stood 2.6 to 4.4 times above their scrambled floors.

The foot and the peak of the pulse are two bins of the brightness trace of the whole field. Every excursion in the paper is the waveform at the peak minus the waveform at the foot. It is referenced to those field-wide moments and never to the extremes of a vessel or of a cut, which would be positive by construction and would lift the floor by as much. Where the phase of a cut does not coincide with that of the field, its foot-to-peak value is smaller than twice its amplitude, so every excursion here is a conservative estimate of the amplitude it stands for.

### Segmentation and hand labelling

The glow around the vessels is estimated once on the mean of the cycle by a morphological opening and subtracted from every phase bin, which changes no displacement and no phase. Pixels are sorted into five categories from parenchyma to lumen by a local-contrast binarisation, the vessel mask is cut into segments at branch points, and the geometry of each segment is measured. The labelling of vessels is the one step done by hand. An operator assembles segments into individual arteries and veins, marks the origin of each vessel (the end at which the blood arrives) and its ends, and gives its type. One centre line is traced from the origin through every branch, and one cut is placed at every pixel of it (Supplementary Fig. 1). A cut is a short line across the vessel that reaches three local radii plus 12 pixels on either side. In the 11 recordings 19 arteries and 19 veins were labelled, and 39 691 cuts returned both channels at every phase bin (18 084 on arteries, 21 607 on veins; 2736 to 5429 per animal).

### The three motions

The preparation moves under the objective, and over the heartbeat this movement is about as large as that of the vessels. It is fitted on background pixels as one similarity transform per phase bin (a slide, a magnification and a rotation) and taken out of the cycle before any vessel is measured. Its foot-to-peak excursion along its own axis is the bulk channel of Fig. 1d, one number per animal. Its magnification (median 0.012 pixels at the radius of the field, 5.7 times its floor) is stored but not drawn.

At each cut the brightness profile is fitted at every phase bin. The fit gives the half-width of the vessel and the transverse position of its centre. The half-width is the mean distance from the centre to the half-maximum crossing on each side, each side against its own baseline. The diameter is twice the half-width. A half-width is blind to translation, so it is independent from flexural motion and bulk translation.

For each labelled vessel with at least 20 cuts, the rigid slide is the single two-dimensional displacement that best explains the transverse displacements of all its cuts in the least-squares sense. The bend is the orthogonal residual, the transverse displacement that varies along the vessel. Over the arteries the rigid slide carries 0.2 % [0.1–0.9] of the transverse power, and the condition number of the fit is 2.5 [1.9–3.6] (largest 8.5). The flexural channel of Fig. 1 is the whole transverse motion of the centre line after the movement of the field is removed, so it is the bend to within that share.

Both channels are smoothed along the vessel with a Gaussian whose full width is the diameter of the vessel, because a bend cannot be sharper than the vessel is wide. The width has a floor of 9 pixels and is 11.3 pixels (30 μm) over the arteries, which is 4.5 % of the spatial period of the bend. The excursion of each vessel is the median over its cuts of the smoothed foot-to-peak change, taken as a magnitude for the flexural channel and signed for the dilatation. The per-animal values of Fig. 1d,e are the medians over the arteries or veins of that animal. The per-wall ratio is the flexural excursion over half the dilatation, because the dilatation is a change of diameter, and it is 3.11 [2.32–4.13] over the 19 arteries. The direction panel (Fig. 2a) bins the arterial cuts of each recording by the direction of their normal in the frame of the removed slide, in 36 bins of 10^°^. A bin is kept only if it holds at least 8 cuts or 2 % of the cuts of the recording, whichever is larger, and the bins are averaged over recordings.

### Phase structure along the artery

The cardiac fundamental of each channel is kept per cut as a complex amplitude. For each labelled artery the longest path from the origin to the farthest marked end is taken, and the flexural fundamental is co-oriented along it. The coherence of the amplitude-weighted fundamental, with magnitudes capped at their 95th percentile, is evaluated over a grid of candidate phase slopes. The spatial period of the flexural wave is 2*π* over the slope that maximises the coherence of the signed, co-oriented fundamental. It is therefore the distance over which the bend repeats with the same sign, which is the bending wavelength *λ* (median 620 μm, interquartile range 504–803 μm, 19 arteries). The energy budget takes *k* = 2*π/λ* from it with no further factor. The confidence interval of the period is the plateau of the coherence curve within the control margin. The diameter on the abscissa of Fig. 2c is the mean lumen diameter along that fitted stretch (median 33.7 μm). The energy budget uses a different estimator, the mean radius along the stretch of the motion fit (median 15.0 μm).

The arrival lag is estimated on the squared fundamental, *z*^2^*/*|*z*|. Squaring turns the sign flips of *π* in the alternation into full turns and so cancels them. What remains is the slow phase drift that an arrival delay leaves along the vessel, doubled, and the conversion divides the doubling out. The search is confined to a declared window of ±30 ms of end-to-end delay, and its control margin is set on that window. The lag is signed positive towards the far end. Per artery it carries a confidence interval of about 5.3 ms, so the test in the text is of its sign over animals. A lag below one frame is resolvable only because the thousands of beats in the averaged cycle vary slightly in duration, and so sample the cycle far more finely than the frame interval.

The camera’s rolling readout delays each sensor line by 9.744 μs times its distance from the sensor’s centre line, which fell at sub-array row 524.5 to 676.5 depending on the line offset of the recording. Across the whole sub-array that is a phase advance of about half a radian at the beat. It cannot be mistaken for a wave that turns by *π* between lobes three times along a stretch, so the readout bears on the lag and not on the alternation. What the readout could leave in the flexural channel, after the vessel’s rigid slide is removed, is a residual that grows with the distance of a cut from the mean sensor line of the vessel and changes sign once, where the vessel crosses that line. Such a residual would be shared by the veins on the same sensor, whose whole flexural channel is 0.032 μm. An artery whose far end lies nearer the centre line than its origin would, however, show its far end lagging even if the physiology were synchronous. The phase along the vessel is therefore analysed after the readout is taken out. Before the fits above, the fundamental of each cut, and of its controls alike, is rotated back by the angular frequency of the beat times the line interval times the distance of the cut in lines from the sensor’s centre line, with the capture offset of each recording. This is the largest delay a centre-out readout can impose.

### Energy budget

Figure 3 shows a model applied to the measurements to estimate how much energy one beat of each motion delivers per metre of artery to three sinks: the viscoelasticity of the wall, the fluid in the perivascular annulus, and the brain tissue outside it. The last two are the two limits of one channel. They are set by a parameter *τ*, the share of the excursion of the artery that the perivascular gap takes up. With the outer wall of the space held (*τ* = 1), the gap is squeezed and the brain does nothing. With that wall free (*τ* = 0), the whole space moves with the artery and the brain carries all of the motion. The fluid losses below are written for the held limit and the tissue losses for the free limit. At intermediate tethering the fluid losses scale as *τ* ^2^ and the tissue losses as (1 − *τ*)^2^.

The artery of lumen radius *R*_1_ lies in an annulus of cerebrospinal fluid that reaches 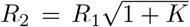, where *K* is the perivascular cross-sectional area over that of the lumen. The gap is *h*_0_ = *R*_2_ − *R*_1_, the mean radius is 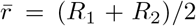, and the wall thickness *h*_*w*_ is a fixed fraction of *R*_1_. The dilatation is the lumen radius rising and falling by an amplitude *A*. The bend is a transverse displacement *ξ* of the axis of the vessel, with curvature *κ* and wavenumber *k* = 2*π/λ*, where *λ* is the bending wavelength. The heart rate *f*_0_ of the recording gives *ω* = 2*πf*_0_ and the period *T* = 1*/f*_0_. From the measurement the model takes, per artery, *R*_1_ (15.0 μm [13.3–16.7]), *f*_0_, *λ* and the mean squares ⟨*A*^2^⟩ and ⟨*ξ*^2^⟩ along the vessel, from which

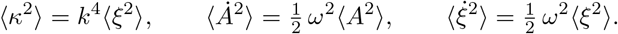

Each foot-to-peak excursion is halved to an amplitude, the dilatation twice because it is a diameter, and squared. For a constant or sinusoidal envelope along the vessel this gives the mean square exactly (root mean squares 43 nm [36–49] for *A* and 146 nm [88–172] for *ξ*). If the bend is a helix that turns out of the imaging plane, which the camera cannot exclude, ⟨*ξ*^2^⟩ is twice the planar reading and every flexural energy doubles exactly. All energies below are per beat and per unit length of vessel.

With *G*^′^ the shear modulus of the wall and tan *δ* its loss tangent, the loss modulus is *G*^′′^ = *G*^′^ tan *δ*, and the viscoelastic losses of the wall are

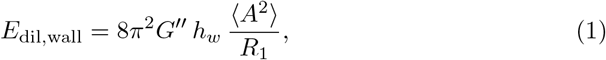

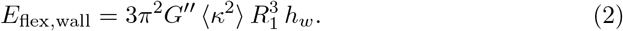

Their ratio is 8/3 times the squared ratio of the hoop strain *A/R*_1_ to the bending strain 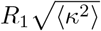, so no material property survives in it.

In the annulus the Womersley number is 0.083 and the Reynolds number 10^−4^, so the flow is quasi-steady Stokes flow. With the CSF viscosity *μ*, the Poiseuille resistance of the annulus per unit length is

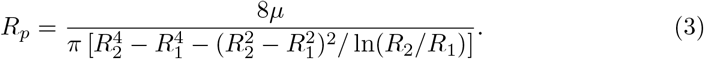

The fluid that the dilatation displaces must leave along the annulus to vents a distance *L* apart, and its loss, averaged along the segment, is

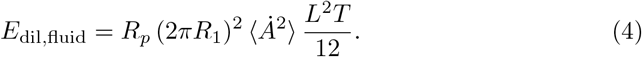

A transverse shift of a cylinder inside a larger one preserves the area between them, so the fluid that the bend displaces is relieved round the annulus and back and forth along the vessel, with zero net axial flux. Its pressure at azimuth *θ* is

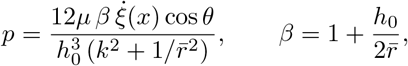

and its loss, the squeeze film relieved by the axial route plus the Couette shear of the wall sliding past the fluid, carries no *L*:

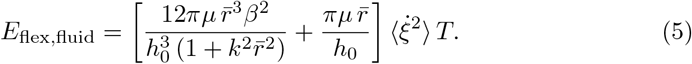

Setting equation (4) equal to equation (5) gives the parity venting length of Fig. 3c per artery. Parity is an equality of two viscous dissipations and not a transport threshold, and the tethering cancels from it because both fluid losses carry *τ* ^2^.

The brain outside is quasi-static and incompressible, with complex shear modulus 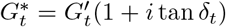 and loss modulus 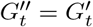 tan *δ*_*t*_. Its losses are

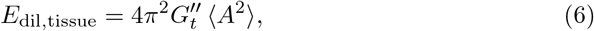

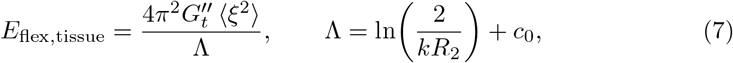

where Λ is the slender-body logarithm that arises because the alternation of the bend cuts its far field off at 1*/k* (Λ = 2.72 [2.37–2.83] over the arteries), and *c*_0_ is a constant of order one. The constants of the model, their sources and the ranges over which they were swept are listed in Supplementary Table 1.

The closed forms were checked against twelve statements written down independently of them, among them exact identities, the classical slit and journal-bearing limits and a direct numerical solution of the linearised Reynolds equation on an axial–azimuthal grid. All twelve hold on five written-down geometries of the measured order.

The model has limits of its own. Its geometry is idealised. The gap is not thin 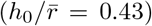, and the wall takes up no room in the annulus although it is about a quarter of the gap. The annulus is concentric and circular, whereas measured pial perivascular spaces are eccentric and often two-lobed [18, 19]. The tissue is unbounded, although a pial artery lies at the cortical surface. The wall is isotropic, with one loss tangent in hoop and along the vessel. An anisotropic wall would put the ratio of its two loss tangents, which is not known for these vessels, into the factor 8/3. Every energy also depends on the smoothing, because both mean squares are taken after it. Before smoothing, the per-cut moment is 14 [5.5–51] times the smoothed one for the bend and 37 [18–63] times for the dilatation, and almost none of the difference is spatially coherent. If it were real motion, every absolute energy would be an underestimate, and the flexure-to-dilatation ratio would fall to 0.40 [0.24–0.90] of what is reported.

The tethering *τ* is the widest sensitivity row, a factor of 37 in the total flexure-to-dilatation ratio, wall included. No other constant moves that ratio by more than a factor of 3 across its swept range (Supplementary Table 1).

### Statistics

The animal is the statistical unit. Per-animal values are medians over the labelled arteries or veins of that animal. Comparisons between channels and between vessel types are two-sided Wilcoxon signed-rank tests over the 11 animals, whose smallest attainable p is 9.8 ×10^−4^. The arrival lag is tested by the sign of the per-animal median over animals (two-sided sign test). The sex comparison is an exact permutation test over the 330 assignments of 4 males among 11 animals (Cliff’s *δ*). Values are medians with the interquartile range (25th and 75th percentiles) in brackets. Medians are used throughout because the quantities of the energy budget span one to two decades across arteries. Correlations are Pearson’s over the 19 arteries, to which the 11 animals contribute one to three each. They are descriptive and are not tests over animals. No correction for multiplicity is applied, and every p is reported where it is drawn.

### Unprocessed video

Supplementary Video 1 is 300 consecutive camera frames (3.0 s) of one recording, not the example recording of Fig. 1, cut to a rectangle of 200 by 300 pixels around one artery and played ten times slower. The frames are not registered, averaged, filtered or magnified.

### Motion-magnified movies

The stills of Fig. 1g and Supplementary Videos 2 and 3 were made by phase-based motion magnification [20]. The averaged cycle is decomposed into a band-passed pyramid (levels 3 to 5, five harmonics of the beat, phase smoothed over 2 pixels of each level). Its phase is amplified by 53.1, 106.2 and 212.5 at the three levels, only within a 3-pixel ring around the segmented vessels, and the cycle is rebuilt. The control movie is the identical magnification of one scrambled-timing control of the same beats, warped by its own fit and shown on the same grey limits.

### Use of large language models

No result or datum in this paper was generated by a model. Claude Code (Anthropic) was used to refine and refactor the analysis code, and Claude Code and Grammarly were used to refine the language of the manuscript.

## Data availability

The eleven results files and the figure data file will be deposited on Zenodo upon acceptance, and the data supplement describing every stored field will be on GitHub with the code. Editors and reviewers can obtain them on request. The raw recordings are available from the corresponding author on reasonable request.

## Code availability

The analysis code (CardiacWallMotion, MIT licence) will be released on GitHub upon acceptance. Editors and reviewers can obtain it on request.

## Acknowledgements

The study was supported by the Lundbeck Foundation (grants R345-2020-1782 and R310-2018-3455).

## Author contributions

M.V.S. performed the experiments; D.D.P. conceived the study and analysed the data; D.D.P., L.Ø. and M.V.S. wrote the manuscript.

