## Supplementary information for "Pial arteries bend with every heartbeat"

**Abstract**

This file holds Supplementary Figures 1 to 4, Supplementary Table 1 (the constants of the energy model, whose equations are in Methods) and the Supplementary References, whose numbering continues from the main paper's reference list (works cited in both are listed again here under their new numbers). Supplementary Videos 1 to 3 are separate files, and their legends are in the Supplementary Information guide.

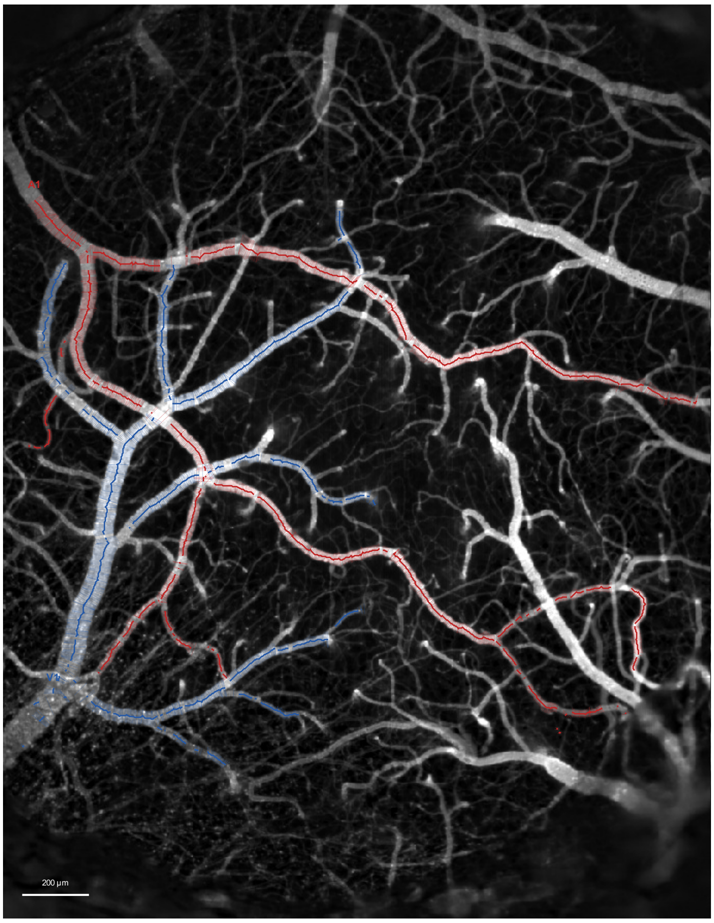


**Supplementary Fig. 1  The labelled vessels of one recording and every accepted cut.** The foot image of the recording is shown as the full common crop of 1034 × 800 pixels, on the stored grey limits, with its two labelled vessel trees and their branches, artery A1 (red) and vein V1 (blue). The centre line of each vessel runs through the centres of its cuts. Every accepted cut is drawn as one semi-transparent segment across the vessel, from wall to wall at its fitted radius, so that crowded cuts appear denser (1972 artery and 1526 vein cuts). Each label is placed at the origin of its tree, which is the end at which the blood arrives. Scale bar, 200 µm. The recording is the same as in Fig. 1a,g.

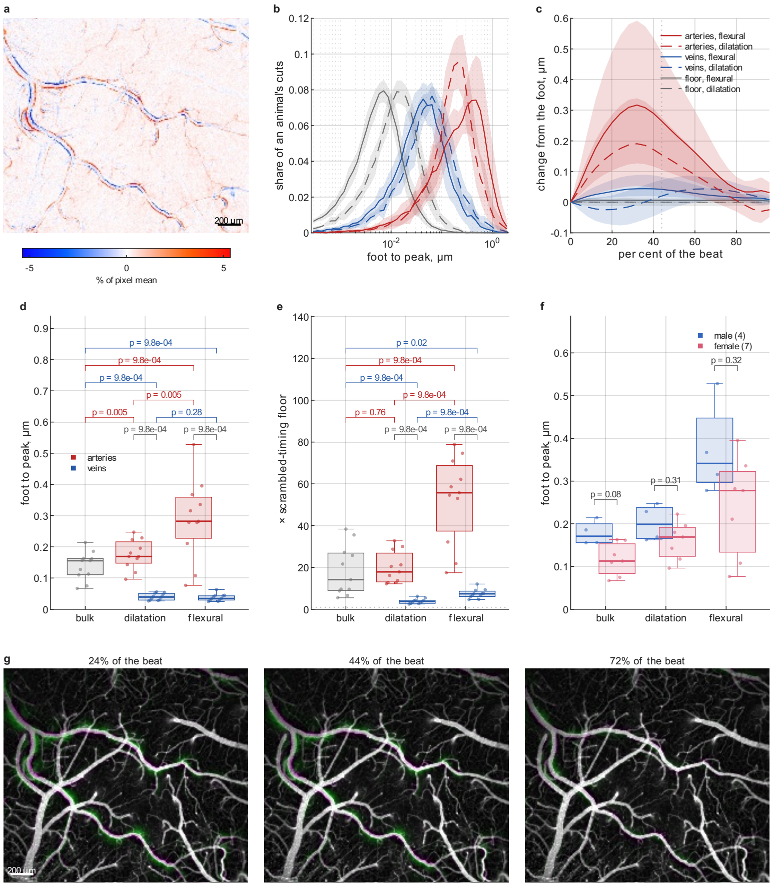


**Supplementary Fig. 2 Figure 1 formed from all six fitted harmonics.** Panels a and g are identical to Fig. 1, and panels b to f are the six-harmonic versions, which serve as a check on the fundamental (Methods). Layout, colours and tests are as in Fig. 1. The per-animal medians are 0.282 µm for the artery flexural excursion, 0.169 µm for the artery dilatation (a change of diameter), 0.035 µm for the vein flexural excursion, 0.039 µm for the vein dilatation and 0.156 µm for the bulk slide, which is unchanged. Over its floor, the artery flexural excursion stands at ×56, the artery dilatation at ×18, the bulk slide at ×14, the vein flexural excursion at ×7.4 and the vein dilatation at ×3.7. Some brackets differ from Fig. 1. In d, the artery bracket of bulk against dilatation gives p = 0.005 and the vein bracket of dilatation against flexural p = 0.28. In e, the artery bracket of bulk against dilatation gives p = 0.76 and the vein bracket of bulk against flexural p = 0.02. In f, the dilatation gives p = 0.31 and the flexural excursion p = 0.32, and the bulk slide is unchanged at p = 0.08.

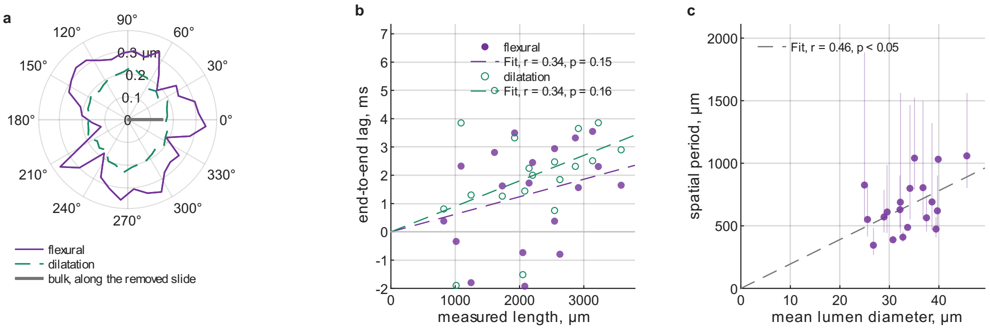


**Supplementary Fig. 3 Figure 2 formed from all six fitted harmonics.** Only panel a differs from Fig. 2, and its outermost tick is at 0.3 µm. Panels b and c are defined on the phase of the cardiac fundamental and are therefore the same in both figures. The layout is as in Fig. 2.

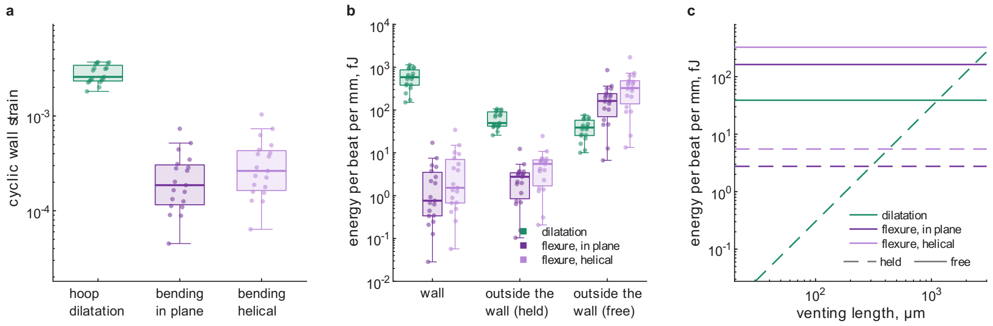


**Supplementary Fig. 4  Figure 3 formed from all six fitted harmonics.** Both excursions are fitted with all six harmonics, so every panel differs slightly from Fig. 3. Layout, colours, line styles and constants are as in Fig. 3. The median hoop strain is 0.258 % and the median bending strain 0.0186 % (0.0263 % as a helix). For the dilatation, the planar bend and the helical bend, the medians in fJ per beat per mm are 584, 0.76 and 1.53 in the wall, 49.6, 2.8 and 5.5 in the held limit, and 38.9, 163 and 325 in the free limit. The parity venting length is 230 µm [167–322] in the plane and 326 µm [236–455] as a helix.

### Supplementary Table

| Quantity | Value | Swept over | Spread | Source |
|---|---|---|---|---|
| perivascular over lumen area, $K$ | 1.4 | 0.5–3 | 2.5 | in vivo mouse pial mean [21, 22] |
| CSF viscosity, $\mu$ | 0.7 mPa s | | | water at 37 °C; human CSF 0.7–1 [23] |
| wall incremental Young modulus | 0.5 MPa | 0.1–1 | 2.8 | rat cerebral artery at 75 mmHg, 0.45–0.62 [24] |
| wall loss tangent, $\tan\delta$ | 0.15 | 0.05–0.3 | 2.1 | 0.1 at 2 Hz to 0.15–0.2 at 18 Hz [25, 26] |
| wall thickness over lumen radius | 0.15 | 0.08–0.25 | 1.6 | pial arteriole wall area and lumen, mouse [27], rat [28] |
| cortical storage shear modulus, $G'_t$ | 2 kPa | 1–3 | 1.0 | mouse, 1.6 kPa at 600 Hz [29]; human 1.2–1.6 at 25–50 Hz [30], 2.7–3.1 at 90 Hz [31] |
| tissue loss tangent, $\tan\delta_t$ | 0.25 | 0.1–0.4 | 1.0 | 0.6–0.7 at 600 Hz [29]; 0.8–0.9 at 90 Hz [31] |
| slender-body constant, $c_0$ | 0.5 | | | order one; enters logarithmically |
| tethering, $\tau$ | 1 | 0–1 | 37.4 | unmeasured |
| venting length, $L$ | 1280 µm | 50–1280 | 1.1 | bifurcation spacing of the labelled arteries of this set [32, 33] |
| penetrating-arteriole band | 50–300 µm | | | |

Supplementary Table 1. The borrowed constants of the energy model of Fig. 3 (Methods): material properties, anatomical lengths and the one choice ($\tau$) that the measurement does not fix. The shear modulus of the wall is one third of the Young modulus, the incompressible limit. Spread is the factor by which the flexure-to-dilatation energy ratio moves across the swept range, with the whole calculation run again at each end (Methods). The two brain rows move nothing, because at the held limit the brain carries none of the motion.
